# An evaluation of the Air Quality Health Index Program on respiratory diseases in Hong Kong: an interrupted time series analysis

**DOI:** 10.1101/529313

**Authors:** Tonya G. Mason, C. Mary Schooling, King Pan Chan, Linwei Tian

**Affiliations:** School of Public Health, Li Ka Shing Faculty of Medicine, The University of Hong Kong, SAR, China

**Keywords:** AQHI, Respiratory diseases, Interrupted time series, Segmented regression

## Abstract

**Background:** On December 30^th^, 2013, the Hong Kong government implemented the Air Quality Health Index (AQHI) to reduce short-term impacts of air pollution on the population. However, whether air quality alert programs, such as the AQHI, reduce morbidity is still questionable. Using a quasi-experimental design, we conducted the first evaluation of the AQHI in Hong Kong focusing on respiratory morbidity.

**Method:** Interrupted time series with Poisson segmented regression from 2010 to 2016 were used to detect any sudden or gradual changes in emergency respiratory hospital admissions, adjusted for air pollutants (NO_2_, SO_2_, PM_10_, O_3_), temperature and humidity, when the AQHI policy was implemented. Findings were validated using three false policy periods. We also assessed changes by specific respiratory diseases (respiratory tract infections (RTI), asthma, chronic obstructive pulmonary disease and pneumonia) and by age.

**Results:** From January 1^st^, 2010-December 31^st^, 2016, 10576.98 deseasonalized, age- and sex-standardized hospital admissions for respiratory diseases occurred in Hong Kong. On implementation of the AQHI, RTI admissions immediately dropped by 14% (relative risk (RR) 0.86 95% confidence interval (CI) 0.76-0.98). In age specific analysis, immediate reductions in hospital admissions, were only apparent in children for RTI (RR 0.84, 95% CI 0.74-0.96) and pneumonia (RR 0.88, 95% CI 0.60-0.96).

**Conclusion:** Hong Kong’s AQHI helped reduced hospital admissions in children, particularly for RTI and pneumonia. To maximize the health benefits of the policy, at risk groups need to be able to follow the behavioral changes recommended by the AQHI index.

## 1. Introduction

Air pollution is a major risk factor for chronic respiratory diseases (World Health Organization 2017a) affecting hundreds of millions of people on a daily basis (World Health Organization 2017b). In 2016 the World Health Organization (WHO) reported air pollution to be responsible for 4.6 million premature deaths with 18% being from respiratory diseases(World Health Organization 2016). Numerous observation studies in both developing and developed countries have shown air pollution associated with hospital admissions for respiratory diseases (Wong et al. 1999; Medina-Ramón, Zanobetti, and Schwartz 2006; Wang et al. 2012; Tao et al. 2014a). Air pollution exacerbates respiratory conditions in “at risk” populations, such as seniors ≥ 65, young children and persons with preexisting respiratory conditions. For example, based on a study conducted in eight European cities, hospital admissions for chronic obstructive pulmonary disease (COPD) plus asthma and total respiratory disease for the elderly (65+), increased by 1.0% and 0.9% for every 10μg /m^3^ increase in particulate matter with aerodynamic diameter ≥10 (PM_10_) (ATKINSON et al. 2001). In addition, hospital admissions for asthma were slightly higher in younger children (0-14) (1.2% change in hospital admission per 10 μg /m^3^ increase in PM_10_) (ATKINSON et al. 2001).

Due to rapid industrialization and economic growth, air pollution has become a major public health issue in China (Xu et al. 2016), with levels of some pollutants well above those generally seen in Western settings(Wong, Tam, and Yu 2002). With an increasing number of vehicles on the road and emissions from industrial activities, Chinese cities are now experiencing very poor air quality, to the point where it reduces visibility(Zhang and Samet 2015). Numerous studies have shown air pollution in major cities, such as Beijing and Hong Kong, is so severe– it has now become a major threat to the human health and well-being. (Wong et al. 1999; Xiong et al. 2015) (Wong et al. 2002; Tao et al. 2014b).

Clearly the long-term solution is inter-governmental action to improve air quality, meanwhile in the short-term, to mitigate public health issues arising from poor air quality, globally, governments have implemented air quality alert programs. These programs advise the public to take precautions when air quality is poor. Despite their popularity, how successful air quality alert programs are at reducing morbidity and mortality is still questionable. Previous evaluations of air quality alert systems have been conducted with some studies finding them to be effective (Neidell 2010; Mullins and Bharadwaj 2014) while others found they have a limited effect on public health(Chen et al. 2018). Moreover, public response to these programs may be culturally or setting specific. Here, we took advantage of the introduction of Hong Kong’s air quality health index (AQHI) alerts (December 30^th^, 2013), aimed at the “at risk” population (seniors ≥65 years, young children, persons with preexisting respiratory and cardiovascular diseases) (EPD 2013a), to assess the effect of such a policy on respiratory diseases. The AQHI advises the public, including “at risk” populations, of the protective measures to take to reduce exposure to poor air quality (CHP 2014)(Table1.). The advice is regularly communicated via local newspapers, radio and television channels on an hourly basis (Environmental Protection Department. The Government of the Hong Kong Special Administrative Region. 2013). AQHI forecasts can also be sent to smartphones and desktop computers by through use of the AQHI “app” (Environmental Protection Department. The Government of the Hong Kong Special Administrative Region. 2013). The majority of the Hong Kong population have access to these services. The AQHI has a 5-point scale low risk, moderate risk, high risk, very high risk and serious (CHP 2014; EPD 2013b) based on a 3 hour-moving average of a combination of air pollutants (nitrogen dioxide (NO_2_), sulfur dioxide (SO_2_), ozone (O_3_) and particulate matter (PM_10_ or PM_2.5_))(EPD 2013b). Here, using a quasi-experimental design, we conducted the first evaluation of the AQHI to assess the effect on respiratory diseases, overall, by subtype and by age in the highly polluted mega-city of Hong Kong.

**Table 1.** Hong Kong Air Quality Health Index (AQHI) health risk categories(Environmental Protection Department The Government of Hong Kong SAR 2013)

| Health Risk Category | AQHI | Health advice |  |  |
| --- | --- | --- | --- | --- |
|  |  | Sensitive Population* | Outdoor Workers | General Public |
| Low | 1-3 | Outdoor activities permitted | Outdoor activities permitted | Ideal for outdoor activities |
| Moderate | 4-6 | Outdoor activities permitted | Outdoor activities permitted | Outdoor activities permitted |
| High | 7 | Reduce outdoor activities and time spent outdoors, especially in areas with heavy traffic | Outdoor activities permitted | Outdoor activities permitted |
| Very High | 8-10 | Reduce outdoor activities and time spent outdoors to a minimum, especially in areas with heavy traffic | Employers should evaluate the risk of keeping their workers outdoor and implement preventive measures that would protect the health of their employees. | Reduce outdoor activities and time spent outdoors, especially in areas with heavy traffic |
| Serious | 10+ | Avoid all outdoor activities and time spent outdoors, particularly in areas with heavy traffic. | Employers should evaluate the risk of keeping their workers outdoor and implement preventive measures that would protect the health of their employees. | Reduce outdoor activities and time spent outdoors to a minimum, especially in areas with heavy traffic |

\*Children and the elderly with existing respiratory conditions

## 2. Materials and Methods

### 2.1 Study population

The study population was the whole population of Hong Kong. Hospital admissions from 2010-16 for all publicly funded hospitals in Hong Kong, accounting for 90% of hospital beds in Hong Kong (Tian et al. 2015; Wong et al. 1999), were obtained from the Hospital Authority for all respiratory diseases. Discharges are routinely coded based on International Classification of Diseases Ninth Revision (ICD9). The population was considered overall, and by age, as children (<20 years), adult (20-<65 years) and elderly (65+years) because of the different vulnerabilities of these age-groups.

### 2.2 Outcomes

Outcomes considered were hospital admissions for respiratory diseases (RDs), including all respiratory diseases (ICD9 460–519), respiratory tract infections (RTIs) (ICD9 460–478), asthma (ICD9 493), chronic obstructive pulmonary disease (COPD) (ICD9 491, 492, 496) and pneumonia (ICD9 480:486). Admissions were classified only on the principal discharge code. As a control outcome, we also considered mortality rates for injuries, poisonings and other external causes (S00-T98), because these would not be expected to respond to the AQHI. These mortality rates were obtained from the Hong Kong Census and Statistics Department from 2010-2015.

### 2.2 Air pollutants and meteorological data

Levels of air pollution (NO_2_, SO_2_, O_3_, PM_10_) were obtained from Hong Kong Environmental Protection Department (HKEPD) from January 2010-December 2016. Hong Kong has 16 fixed air pollution monitoring stations: 3 are roadside stations and the other 13 are general stations. Three of the general stations were excluded due to extensive missing data. The 10 monitoring stations are representative of Hong Kong’s population exposure because the majority of Hong Kong’s population lives within 5 kilometers of these stations (Huang, Leung, and Schooling 2017). We obtained monthly mean pollution concentrations by averaging each pollution concentration across the 10-monitoring station. Monthly mean temperature and relative humidity were obtained from the Hong Kong Observatory from 2010-2016.

### 2.4 Study Design and Statistical Analyses

To evaluate the effectiveness of the introduction of the AQHI in December 2015, we used an interrupted time series study with a Poisson segmented regression analysis from January 2010-December 2016 to identify sudden and gradual changes in age-standardized hospital admissions for respiratory diseases (Nistal-Nuño 2017). Hospital admissions were age-and sex-standardized based on the WHO standard population. The seasonal trend was adjusted for by using the seasonal trend decomposition loess (STL) procedure, to retrieve the deseasonalized data prior to carrying out the segmented regression models (Cleveland et al. 1990; Jassim, Coskuner, and Munir 2018).

Deseasonalized, age-and sex-standardized respiratory hospitalizations (Y*t*) were regressed on time (see equation below). The mean number of hospitalizations at the beginning of the study period, baseline period, are represented as (β_0_). Trends over time are captured by (β_1_). The AQHI intervention is represented as (β_2_), it takes the value of 0 for the pre-intervention period, and 1 for post intervention. Gradual changes were detected by including an interaction term between the policy and time (β_3_). All other covariates adjusted for, i.e., air pollutants, temperature and humidity, are captured by β_4_. [eq]

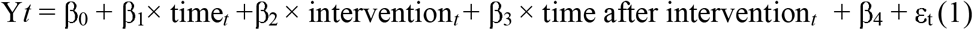

Residuals were plotted to check for over dispersion.

Model validity was tested by implemented three false policy periods 6 months, 12 months and 24 months before the intervention (Stallings-Smith et al. 2013).

All statistical analysis were conducted using the software package R 3.4.2.

#### Ethics

For this entire study aggregated data were used, thus ethical approval is not required.

## 3. Results

The monthly mean respiratory hospital admissions overall and by disease are displayed in Table 2. Monthly mean hospital admissions for all respiratory diseases were 125.9 with a standard deviation (SD) of 23.83. For individual respiratory diseases, pneumonia had the highest hospital admission (mean: 48.12; SD: 8.28) followed by RTIs (mean: 26.18; SD: 5.42). On average, there were 23.70 (SD: 4.83) and 7.85 (SD: 1.37) hospital admissions per month, for COPD and asthma. Influenza had mean monthly hospital admissions of 23.45(SD: 4.95). NO2, SO2, PM10 and O3 had monthly mean concentrations of 49.12 (SD:10.15), 11.27 (SD: 2.64), 41.88 (SD: 16.84), 42.04 (SD: 12.73), respectively. Overall from January 2010 to December 2016 10576.98 hospital admission for respiratory diseases occurred.

**Table 2.** Summary statistics for hospital admissions (counts per day) for respiratory diseases and covariates for the entire study period (2010-2016).

|  | Mean | SD | Min | 25th | Med | 75th | Max |
| --- | --- | --- | --- | --- | --- | --- | --- |
| **Predicted variables** |  |  |  |  |  |  |  |
| Respiratory diseases | 125.92 | 23.83 | 90.37 | 106.92 | 121.22 | 137.21 | 191.50 |
| RTI | 26.18 | 5.42 | 17.75 | 22.55 | 24.94 | 28.60 | 41.65 |
| COPD | 23.70 | 4.83 | 15.53 | 19.51 | 23.01 | 27.14 | 36.65 |
| Asthma | 7.85 | 1.37 | 5.01 | 6.93 | 7.71 | 8.48 | 11.26 |
| Pneumonia | 48.12 | 8.28 | 35.84 | 42.34 | 47.15 | 51.73 | 70.93 |
| **Covariates†** |  |  |  |  |  |  |  |
| NO<sub>2</sub>(ug/m<sup>3</sup>) | 49.12 | 10.15 | 28.61 | 41.70 | 48.90 | 55.23 | 71.53 |
| SO<sub>2</sub>(ug/m<sup>3</sup>) | 11.27 | 2.64 | 6.68 | 9.54 | 10.79 | 12.69 | 19.82 |
| PM<sub>10</sub>(ug/m<sup>3</sup>) | 41.88 | 16.84 | 16.5 | 27.66 | 39.84 | 53.84 | 84.21 |
| O<sub>3</sub>(ug/m<sup>3</sup>) | 42.04 | 12.73 | 18.54 | 32.48 | 41.58 | 49.38 | 84.12 |

RTI = Respiratory tract infection, COPD= Chronic obstructive pulmonary disease.

Table 3 summarizes the immediate and gradual effects of the AQHI policy on respiratory diseases, illustrated in figure 1. Following the policy an immediate decrease was observed for RTIs, which fell 14% (relative risk (RR) 0.86 95% confidence interval (CI) 0.76-0.98), followed by a static trend (RR 1.00 95% CI 0.99-1.00). In contrast, no immediate or gradual decreases were observed in hospital admissions for all respiratory diseases, COPD, asthma or pneumonia after implementation of the policy, although the estimate for overall respiratory admissions was RR 0.92, 95% CI 0.81-1.04, and for and pneumonia was RR 0.88, 95% CI 0.88-1.20. No gradual post policy change was evident for any disease considered.

**Table 3.** Approximation of immediate and gradual changes of respiratory diseases after implementation of the AQHI policy in a multivariate analysis†.

| Diseases | Immediate Effects | Gradual Effects |
| --- | --- | --- |
|  | RR (95% CI) | RR (95% CI) |
| All Respiratory diseases | 0.92 (0.81-1.04) | 1.00 (1.00-1.01) |
| Respiratory tract infection | **0.86 (0.76-0.98)** | 1.00 (0.99-1.01) |
| COPD | 0.97 (0.90-1.05) | 1.00 (1.00-1.01) |
| Asthma | 1.03 (0.91-1.17) | 1.00 (1.00-1.01) |
| Pneumonia | 0.88 (0.77-1.00) | 1.01 (1.00-1.01) |
| Injuries and External causes | 1.36 (0.92-2.02) | 0.96 (0.94-0.99) |

†Adjusted for seasonality, time trend, SO<sub>2</sub>, NO<sub>2</sub>, PM<sub>10</sub>, O<sub>3</sub>.
‡Age standardized based on the World Standard Population.
RTI = Respiratory tract infection, COPD= Chronic obstructive pulmonary disease.

**Fig. 1.**
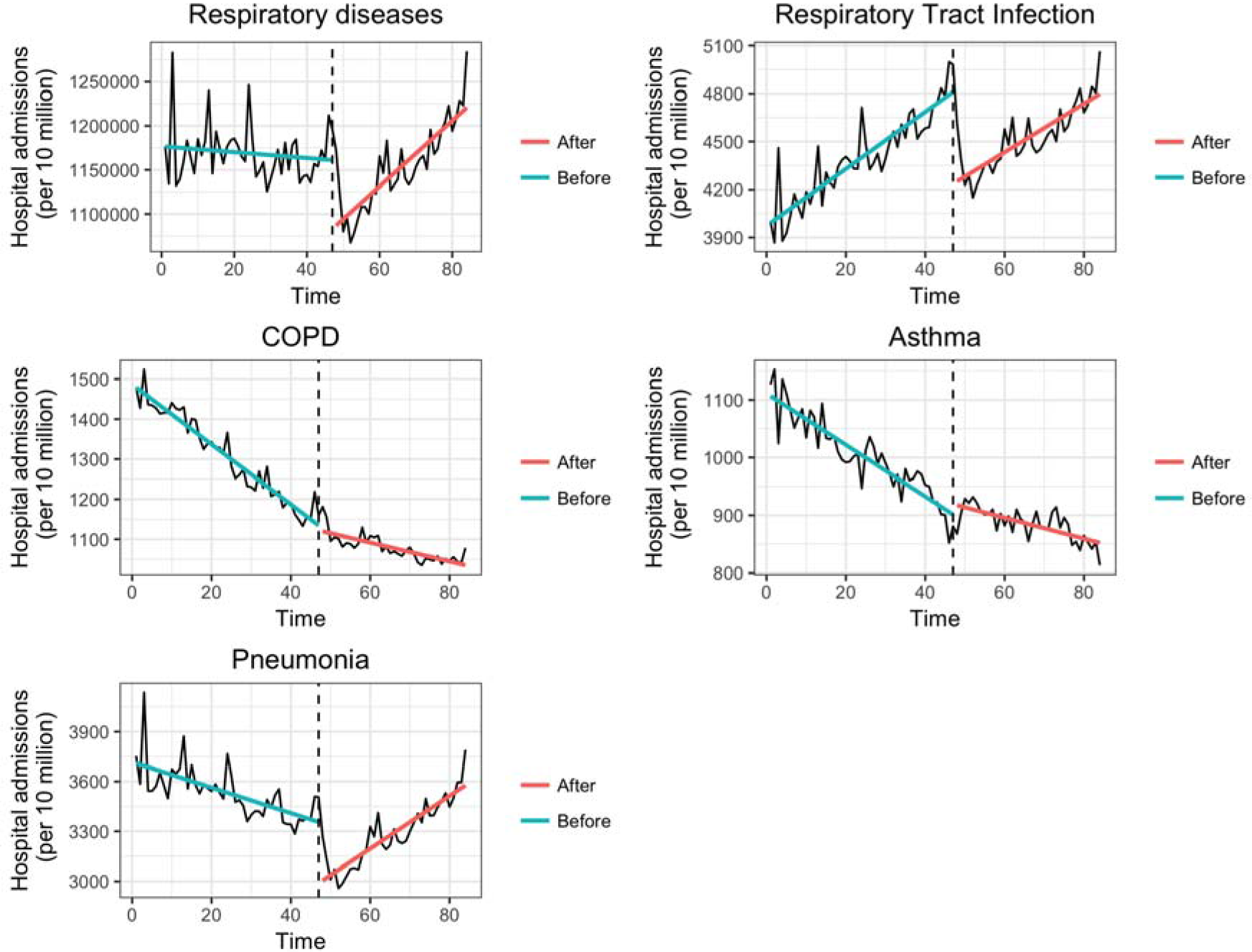
Time series plot of monthly mean, age- and sex-standardised emergency hospital admissions: adjusted for; seasonality, air pollutants and time trend, for all and specific respiratory diseases in Hong Kong, 2010-2016.

Age stratified estimates are displayed in table 4. Post policy the only significant immediate decrease was detected in children (<20 years) for RTI and pneumonia hospital admissions. In children RTI dropped by 16% (RR 0.84, 95% CI 0.74-0.96) and pneumonia by 24% (RR 0.88, 95% CI 0.60-0.96) respectively (see supplementary Figures 3 & 4). An increasing annual trend was observed after the policy for both diseases. In contrast, for all other diseases and age-groups considered no immediate decrease or changes in annual trend were observed after the AQHI policy was implemented. A non-significant reduction in respiratory diseases was only observed in children (<20 years) immediately after the policy implementation (RR 0.86, 95% CI 0.74-1.00) (Figure 2).

**Table 4.** Approximation of immediate and gradual changes for respiratory diseases by age category^‡^ post AQHI policy in a multivariate analysis†.

|  | >20 Years |  | 20-65 Years |  | < 65 Years |  |
| --- | --- | --- | --- | --- | --- | --- |
| Diseases | Immediate Effects | Gradual Effects | Immediate Effects | Gradual Effects | Immediate Effects | Gradual Effects |
|  | RR (95% CI) | RR (95% CI) | RR (95% CI) | RR (95% CI) | RR (95% CI) | RR (95% CI) |
| All Respiratory | 0.86 (0.74-1.00) | 1.00 (1.00-1.01) | 0.99 (0.88-1.11) | 1.01 (1.00-1.01) | 1.01 (0.91-1.13) | 1.00(1.00-1.01) |
| RTI | **0.84 (0.74-0.96)** | 1.00 (0.99-1.01) | 1.01 (0.86-1.19) | 1.00 (0.99-1.01) | 0.98 (0.80-1.21) | 1.00 (0.99-1.00) |
| COPD | 0.46 (0.15-1.37) | 0.97 (0.92-1.03) | 1.04 (0.95-1.13) | 1.00 (1.00-1.01) | 0.96 (0.88-1.04) | 1.00 (1.00-1.01) |
| Asthma | 1.07 (0.90-1.29) | 1.00 (0.99-1.01) | 0.94 (0.85-1.03) | 1.00 (1.00-1.01) | 0.99 (0.88-1.12) | 1.00 (0.99-1.00) |
| Pneumonia | **0.76 (0.60-0.96)** | 1.01 (1.00-1.02) | 0.90 (0.79-1.04) | 1.01 (1.01-1.02) | 1.00 (0.92-1.09) | 1.00 (1.00-1.01) |

†Adjusted for seasonality, time trend, SO<sub>2</sub>, NO<sub>2</sub>, PM<sub>10</sub>, O<sub>3</sub>.
‡Age standardized based on the World Standard Population.
RTI = Respiratory tract infection, COPD= Chronic obstructive pulmonary disease.

**Fig. 2.**
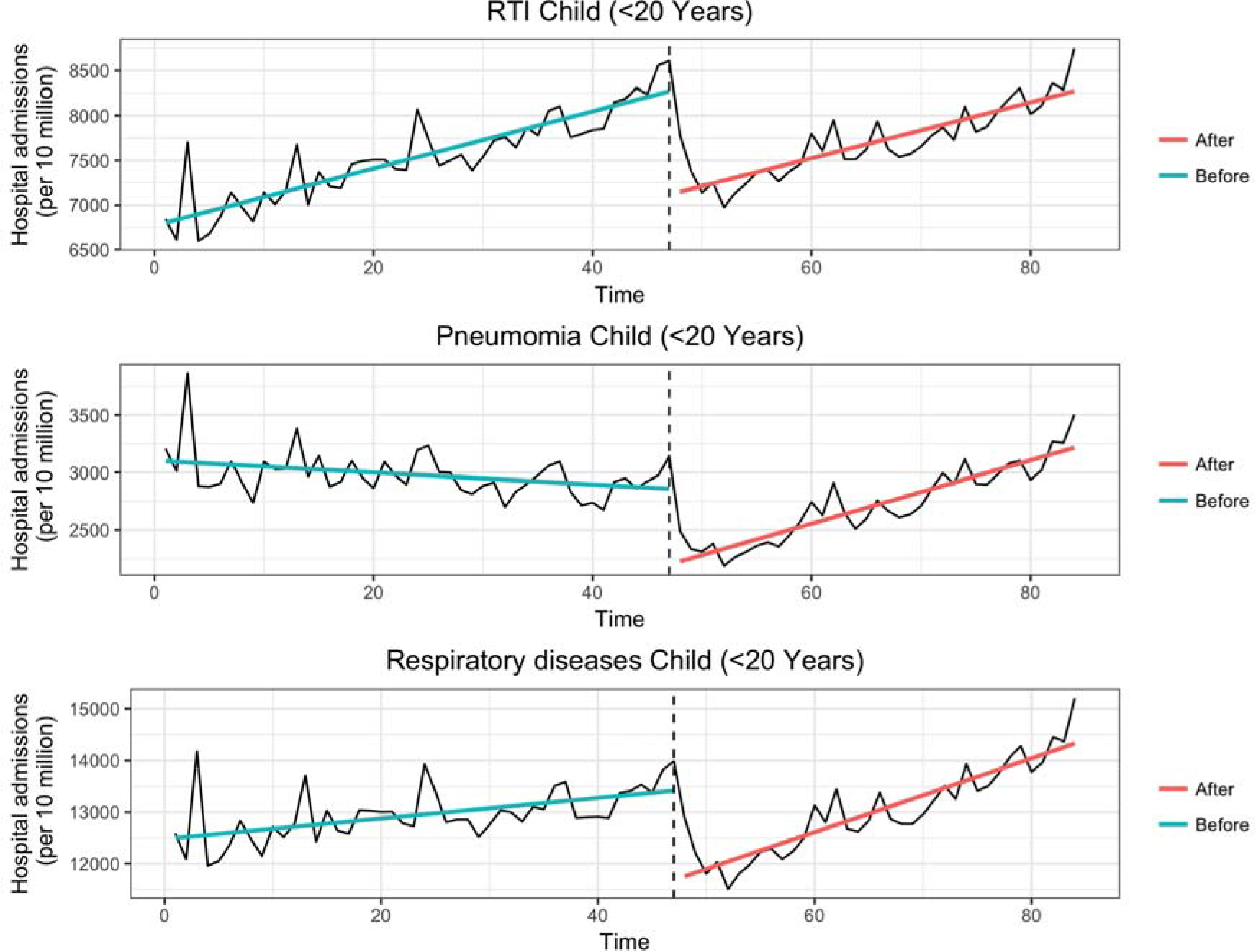
Time series plot of monthly mean, age- and sex-standardised hospital admissions, adjusted for; seasonality, air pollutants and time trend, for RTI, Pneumonia and respiratory disease, in Hong Kong, 2010-2016.

Using the false policy implementation dates of 6, 12 and 24 months before the policy, revealed no significant immediate or gradual decreases in hospital admissions for most diseases considered, except for COPD at a false period of 12 months (see supplementary Table 3). Repeating the analysis for the control outcome of deaths from injuries and external causes gave no immediate changes when the AQHI policy was implemented (see supplementary Figure 5).

## 4. Discussion

Hong Kong’s AQHI policy implementation was associated with an immediate reduction in RTI hospitalizations, in particular for children but did not clearly affect respiratory disease hospitalizations overall, or hospitalizations for COPD, asthma and pneumonia. Our findings are somewhat consistent with other intervention studies evaluating air quality alert impacts in Canada (Chen et al. 2018) and the United States (US) (Neidell 2010). Like our study, the study conducted in Canada did not observe any significant immediate or gradual declining changes for asthma or COPD hospital admissions after the implementation of the AQHI. The US study (Neidell 2010) focused on the effect air quality alerts pertaining to ozone on outdoor activities but not health effects. They found children more responsive to air quality warnings (25% decrease in zoo attendance).

Our findings add to the literature concerning the impact air quality alerts on respiratory diseases, particularly for RTI where published evidence is limited. Air pollution is strongly associated with RTIs, particularly in children (Li et al. 2018; Darrow et al. 2014; Bono et al. 2016; Ghosh et al. 2012). Prior work done by researchers in Rome has demonstrated an association between NO_2_ and O_3_ and hospital admissions for respiratory tract infections in children(Fusco et al. 2001). Out of all respiratory diseases, NO_2_ had a stronger effect on respiratory tract infections on same day lag period (lag0,4.0%increase) (Fusco et al. 2001). As such, any decline in RTI with the implementation of AQHI warnings is likely to be the result of caretakers—likely women, keeping children indoors in response to the AQHI warnings. Woman have a greater concern for the effects of air pollution on health, than men (Flynn, Slovic, and Mertz 1994; Lu 2015). The sudden drop in pneumonia hospital admissions for children is consistent with the fall in RTI admissions. Air pollution is known to be associated with pneumonia hospital admissions in children (Negrisoli and Nascimento 2013; Souza and Nascimento 2016; Lv et al. 2017)

Similar to other cities in the world (UK, Canada, U.S.A); Hong Kong’s AQHI policy alerts the public to risk and thereby promotes behavioral changes for “at risk populations” (Radisic and Newbold 2016). As such, health effects of the policy only occur if the population at risk changes its behavior in response to the information as well as the associated lower exposure reducing disease. A qualitative study done by Radisic and Newbold in 2016 concluded that proper adaption to the AQHI depended on knowledge about the AQHI and characteristics of the policy; and perception about air quality and health. People less educated about the AQHI were less likely to adopt the behavioral changes suggested by the policy, which may include the elderly (Radisic and Newbold 2016). Older people are most vulnerable to COPD and pneumonia but they may not respond immediately to new information (Radisic and Newbold 2016). This consideration may be particularly relevant in Hong Kong where a substantial proportion of older people are illiterate, and so may find it harder to access the AQHI. As for hospital admissions for asthma, these are less clearly linked to air pollution, given the mixed findings from previous studies (Schwartz 1996; Dab et al. 1996; Katsouyanni et al. 1997). Past studies have shown a stronger association of grass pollen (released before and during thunderstorms) with asthma hospital admissions than air pollution(Celenza et al. 1996; Davidson et al. 1996) (Gouveia and Fletcher 2000). Furthermore, emergency hospital admission in Hong Kong consists of severe cases of asthma attacks. The less severe cases are usually advised to be treated at private or public clinics and never recorded under hospital admissions dataset because more likely the patient leaves immediately after medical care (Hospital Authority 2018). Asthma results for our study aligns with a recent air quality intervention study done in Canada that reported a significant drop in emergency admissions for asthma but none for hospital admissions (Chen et al. 2018). Recently, in Hong Kong air pollution is not clearly linked to hospital admissions for asthma.

To the best of our knowledge, this study is the first study to document the AQHI effect on respiratory diseases in Hong Kong. Another major key strength of our study, is the study design. The use of the interrupted time series analysis is known to be a very powerful evaluative design when randomized control studies are not feasible (Lopez Bernal, Cummins, and Gasparrini 2016). The design allowed us to account for secular trends (Penfold and Zhang 2013). The use of a negative control outcome is also a strength of the study because it can be used to detect unknown time varying confounders, and thereby strengthens the validity of our findings. Our study also sheds some light on the potential effects AQHI may have on other respiratory diseases such as RTIs for which there is limited published evidence. Like all studies, we also have some weaknesses. Firstly, ITS cannot be used to make causal inferences on an individual level due to the fact that we used aggregated data (hospital admissions) instead of individual data. However, the target of policy is population health, is assessed here. Secondly, our study included only hospital admissions and lacked numbers from other outpatient settings such as physician visits. However, in these circumstances it is particularly notable that we found an effect on RTIs. Thirdly, our study did not include a sex specific analysis, which may clarify whether there are sex-specific responses to health information. Future studies should assess sex-specific effects of the policy. Lastly, response to an AQHI depends on the public both accessing the relevant information and being able to act on it. We do not have information about the level of awareness of the AQHI in the general population, nor do we know whether everybody was able to respond to the AQHI warnings. For example, people working in outdoor jobs might not be able to take the recommended action. Given, the effectiveness of the AQHI in our setting further research on awareness might inform the implementation of similar policies. Consideration should also be given to the feasibility of the population responding to AQHI warnings

## 5. Conclusion

The implementation of Hong Kong’s AQHI policy resulted in health benefits specifically a reduction in RTI and pneumonia hospital admissions amongst children. To maximize the health benefits of the policy, at risk populations need to be aware on the information, free to take the suggested actions and willing to do so. This study provides valuable information about the effectiveness of the AQHI to policy makers and may inform the decision to take further action if needed.

## 6. Acknowledgements

We thank the Center for Health Protection (CHP) report for supplying the health data and the Environmental Protection Department of the Hong Kong Special Administrative Region for providing the air monitoring data for this study.

## References

Atkinson, Richard W., H. ROSS Anderson, Jordi Sunyer, JON Ayres, Michela Baccini, Judith M. Vonk, Azzedine Boumghar, et al. 2001. “Acute Effects of Particulate Air Pollution on Respiratory Admissions.” American Journal of Respiratory and Critical Care Medicine 164 (10). American Thoracic Society New York, NY: 1860–66. doi:10.1164/ajrccm.164.10.2010138.

Celenza, A, J Fothergill, E Kupek, and R J Shaw. 1996. “Thunderstorm Associated Asthma: A Detailed Analysis of Environmental Factors.” BMJ (Clinical Research Ed.) 312 (7031). British Medical Journal Publishing Group: 604–7. doi:10.1136/BMJ.312.7031.604.

Chen, Hong, Qiongsi Li, Jay S Kaufman, Jun Wang, Ray Copes, Yushan Su, and Tarik Benmarhnia. 2018. “Effect of Air Quality Alerts on Human Health: A Regression Discontinuity Analysis in Toronto, Canada.” The Lancet Planetary Health 2 (1). Elsevier: e19–26. doi:10.1016/S2542-5196(17)30185-7.

CHP. 2014. “Air Quality Health Index Has Come into Effect.” https://www.chp.gov.hk/files/pdf/aqhi_eng_final_20140103.pdf.

Cleveland, Robert B, William S Cleveland, Jean E McRae, and Irma Terpenning. 1990. “STL: A Seasonal-Trend Decomposition Procedure Based on Loess.” Journal of Official Statistics 6 (1): 3–73. http://www.nniiem.ru/file/news/2016/stl-statistical-model.pdf.

Dab, W, S Medina, P Quénel, Y Le Moullec, A Le Tertre, B Thelot, C Monteil, et al. 1996. “Short Term Respiratory Health Effects of Ambient Air Pollution: Results of the APHEA Project in Paris.” Journal of Epidemiology and Community Health 50 Suppl 1 (Suppl 1). BMJ Publishing Group Ltd: s42–6. doi:10.1136/JECH.50.SUPPL_1.S42.

Davidson, A C, J Emberlin, A D Cook, K M Venables, Ruth Brown, Alice Findlay, Miriam Harris, et al. 1996. “A Major Outbreak of Asthma Associated with a Thunderstorm: Experience of Accident and Emergency Departments and Patients’ Characteristics. Thames Regions Accident and Emergency Trainees Association.” BMJ (Clinical Research Ed.) 312 (7031). British Medical Journal Publishing Group: 601–4. doi:10.1136/BMJ.312.7031.601.

Environmental Protection Department. The Government of the Hong Kong Special Administrative Region. 2013. “How Can I Get the AQHI Information? - Page 6.” http://www.aqhi.gov.hk/en/what-is-aqhi/about-aqhia37a.html?showall=&start=5.

EPD. 2013a. “EPD - About AQHI.” http://www.aqhi.gov.hk/en/what-is-aqhi/about-aqhi.html.

EPD. 2013b. “EPD - About AQHI.”

Flynn, James, Paul Slovic, and C. K. Mertz. 1994. “Gender, Race, and Perception of Environmental Health Risks.” Risk Analysis 14 (6). Blackwell Publishing Ltd: 1101–8. doi:10.1111/j.1539-6924.1994.tb00082.x.

Fusco, D, F Forastiere, P Michelozzi, T Spadea, B Ostro, M Arcà, and C A Perucci. 2001. “Air Pollution and Hospital Admissions for Respiratory Conditions in Rome, Italy.” The European Respiratory Journal 17 (6). European Respiratory Society: 1143–50. http://www.ncbi.nlm.nih.gov/pubmed/11491157.

Gouveia, Nelson, and Tony Fletcher. 2000. “Respiratory Diseases in Children and Outdoor Air Pollution in São Paulo, Brazil: A Time Series Analysis.” Occup Environ Med 57: 477–83. doi:10.1136/oem.57.7.477.

Hospital Authority. 2018. “Accident and Emergency (A&E).” http://www.ha.org.hk/visitor/ha_visitor_index.asp?Content_ID=10051&Lang=ENG&Dimension=100&Ver=HTML.

Huang, Jian V, Gabriel M Leung, and C Mary Schooling. 2017. “The Association of Air Pollution With Pubertal Development: Evidence From Hong Kong’s “ Children of 1997 ” Birth Cohort.” American Journal of Epidemiology 185 (10). doi:10.1093/aje/kww200.

Jassim, Majeed S., Gulnur Coskuner, and Said Munir. 2018. “Temporal Analysis of Air Pollution and Its Relationship with Meteorological Parameters in Bahrain, 2006-2012.” Arabian Journal of Geosciences 11 (3). Springer Berlin Heidelberg: 62. doi:10.1007/s12517-018-3403-z.

Katsouyanni, K, G Touloumi, C Spix, J Schwartz, F Balducci, S Medina, G Rossi, et al. 1997. “Short-Term Effects of Ambient Sulphur Dioxide and Particulate Matter on Mortality in 12 European Cities: Results from Time Series Data from the APHEA Project. Air Pollution and Health: A European Approach.” Bmj 314 (7095): 1658–63.

Lopez Bernal, James, Steven Cummins, and Antonio Gasparrini. 2016. “Interrupted Time Series Regression for the Evaluation of Public Health Interventions: A Tutorial.” International Journal of Epidemiology 46 (1). Oxford University Press: 348. doi:10.1093/ije/dyw098.

Lu, Yuanan. 2015. “Research Residents’ Perception of Air Quality, Pollution Sources, and Air Pollution Control in Nanchang, China ArticleHistory.” Atmospheric Pollution Research 6: 835–41. doi:10.5094/APR.2015.092.

Lv, Chenguang, Xianfeng Wang, Na Pang, Lanzhong Wang, Yuping Wang, Tengfei Xu, Yu Zhang, Tianran Zhou, and Wei Li. 2017. “The Impact of Airborne Particulate Matter on Pediatric Hospital Admissions for Pneumonia among Children in Jinan, China: A Case-Crossover Study The Impact of Airborne Particulate Matter on Pediatric Ho.” Journal of the Air & Waste Management Association 67 (6): 669–76. doi:10.1080/10962247.2016.1265026.

Medina-Ramón, Mercedes, Antonella Zanobetti, and Joel Schwartz. 2006. “The Effect of Ozone and PM10 on Hospital Admissions for Pneumonia and Chronic Obstructive Pulmonary Disease: A National Multicity Study.” American Journal of Epidemiology 163 (6). Oxford University Press: 579–88. doi:10.1093/aje/kwj078.

Mullins, Jamie, and Prashant Bharadwaj. 2014. “EFFECTS OF SHORT-TERM MEASURES TO CURB AIR POLLUTION: EVIDENCE FROM SANTIAGO, CHILE.” doi:10.1093/ajae/aau081.

Negrisoli, Juliana, and Luiz Fernando C Nascimento. 2013. “Atmospheric Pollutants and Hospital Admissions Due to Pneumonia in Children.” Revista Paulista de PediatriaJ: Orgao Oficial Da Sociedade de Pediatria de Sao Paulo 31 (4). Sociedade De Pediatria De Sao Paulo: 501–6. doi:10.1590/S0103-05822013000400013.

Neidell, Matthew. 2010. “Air Quality Warnings and Outdoor Activities: Evidence from Southern California Using a Regression Discontinuity Design.” Journal of Epidemiology and Community Health 64 (10). BMJ Publishing Group Ltd: 921–26. doi:10.1136/jech.2008.081489.

Nistal-Nuño, Beatriz. 2017. “Segmented Regression Analysis of Interrupted Time Series Data to Assess Outcomes of a South American Road Traffic Alcohol Policy Change.” Public Health 150 (September). W.B. Saunders: 51–59. doi:10.1016/J.PUHE.2017.04.025.

Penfold, Robert B, and Fang Zhang. 2013. “Use of Interrupted Time Series Analysis in Evaluating Health Care Quality Improvements.” ACAP 13: S38–44. doi:10.1016/j.acap.2013.08.002.

Radisic, Sally, and K Bruce Newbold. 2016. “Factors Influencing Health Care and Service Providers’ and Their Respective “at Risk” Populations’ Adoption of the Air Quality Health Index (AQHI): A Qualitative Study.” BMC Health Services Research 16 (March). BioMed Central: 107. doi:10.1186/s12913-016-1355-0.

Schwartz, Joel. 1996. “Air Pollution and Hospital Admissions for Respiratory Disease.” Source: Epidemiology 7 (1): 20–28. http://www.jstor.org/stable/3702752.

Souza, Laís Salgado Vieira de, and Luiz Fernando Costa Nascimento. 2016. “Air Pollutants and Hospital Admission Due to Pneumonia in Children: A Time Series Analysis.” Revista Da Associação Médica Brasileira 62 (2): 151–56. doi:10.1590/1806-9282.62.02.151.

Stallings-Smith, Sericea, Ariana Zeka, Pat Goodman, Zubair Kabir, and Luke Clancy. 2013. “Reductions in Cardiovascular, Cerebrovascular, and Respiratory Mortality Following the National Irish Smoking Ban: Interrupted Time-Series Analysis.” Edited by Lamberto Manzoli. PLoS ONE 8 (4). Public Library of Science: e62063. doi:10.1371/journal.pone.0062063.

Tao, Yan, Shengquan Mi, Shuhong Zhou, Shigong Wang, and Xiaoyun Xie. 2014a. “Air Pollution and Hospital Admissions for Respiratory Diseases in Lanzhou, China.” Environmental Pollution 185: 196–201. doi:10.1016/j.envpol.2013.10.035.

Tao, Yan, Shengquan Mi, Shuhong Zhou, Shigong Wang, and Xiaoyun Xie. 2014b. “Air Pollution and Hospital Admissions for Respiratory Diseases in Lanzhou, China.” Environmental Pollution 185: 196–201. doi:10.1016/j.envpol.2013.10.035.

Tian, Linwei, Hong Qiu, Vivian C Pun, Kin-Fai Ho, Chi Sing Chan, and Ignatius T S Yu. 2015. “Carbon Monoxide and Stroke: A Time Series Study of Ambient Air Pollution and Emergency Hospitalizations ☆,☆☆,★,★★.” International Journal of Cardiology 201: 4–9. doi:10.1016/j.ijcard.2015.07.099.

Wang, Minzhen, Shan Zheng, Shigong Wang, Yan Tao, and Kezheng Shang. 2012. “[A Time-Series Study on the Relationship between Gaseous Air Pollutants and Daily Hospitalization of Respiratory Disease in Lanzhou City].” Wei Sheng Yan Jiu = Journal of Hygiene Research 41 (5): 771–75. http://www.ncbi.nlm.nih.gov/pubmed/23213692.

Wong, T W, T S Lau, T S Yu, A Neller, S L Wong, W Tam, and S W Pang. 1999. “Air Pollution and Hospital Admissions for Respiratory and Cardiovascular Diseases in Hong Kong.” Occup Environ Med 56 (10): 679–83.

Wong, T W, W S Tam, and T S Yu. 2002. “Associations between Daily Mortalities from Respiratory and Cardiovascular Diseases and Air Pollution in Hong Kong, China.” Occupational and Environmental Medicine 59 (1): 30–35. doi:10.1136/oem.59.1.30.

Wong, T W, W S Tam, T S Yu, and A H S Wong. 2002. “Associations between Daily Mortalities from Respiratory and Cardiovascular Diseases and Air Pollution in Hong Kong, China.” Occup Environ Med 59: 30–35. http://oem.bmj.com/content/oemed/59/1/30.full.pdf.

World Health Organization. 2016. “WHO | Ambient (Outdoor) Air Quality and Health.” Who. http://www.who.int/news-room/fact-sheets/detail/ambient-(outdoor)-air-quality-and-health.

World Health Organization. 2017a. “WHO | Air Pollution.” WHO. World Health Organization. http://www.who.int/ceh/risks/cehair/en/.

World Health Organization. 2017b. “WHO | Ambient Air Quality.” Who. World Health Organization. http://www.who.int/phe/health_topics/outdoorair/en/.

Xiong, Qiulin, Wenji Zhao, Zhaoning Gong, Wenhui Zhao, and Tao Tang. 2015. “Fine Particulate Matter Pollution and Hospital Admissions for Respiratory Diseases in Beijing, China.” International Journal of Environmental Research and Public Health 12 (9). Multidisciplinary Digital Publishing Institute: 11880–92. doi:10.3390/ijerph120911880.

Xu, Qin, Xia Li, Shuo Wang, Chao Wang, Fangfang Huang, Qi Gao, Lijuan Wu, et al. 2016. “Fine Particulate Air Pollution and Hospital Emergency Room Visits for Respiratory Disease in Urban Areas in Beijing, China, in 2013.” PloS One 11 (4). Public Library of Science: e0153099. doi:10.1371/journal.pone.0153099.

Zhang, Junfeng Jim, and Jonathan M Samet. 2015. “Chinese Haze versus Western Smog: Lessons Learned.” Journal of Thoracic Disease 7 (1). AME Publications: 3–13. doi:10.3978/j.issn.2072-1439.2014.12.06.

