## Supplementary material for "An evaluation of the Air Quality Health Index Program on respiratory diseases in Hong Kong: an interrupted time series analysis"

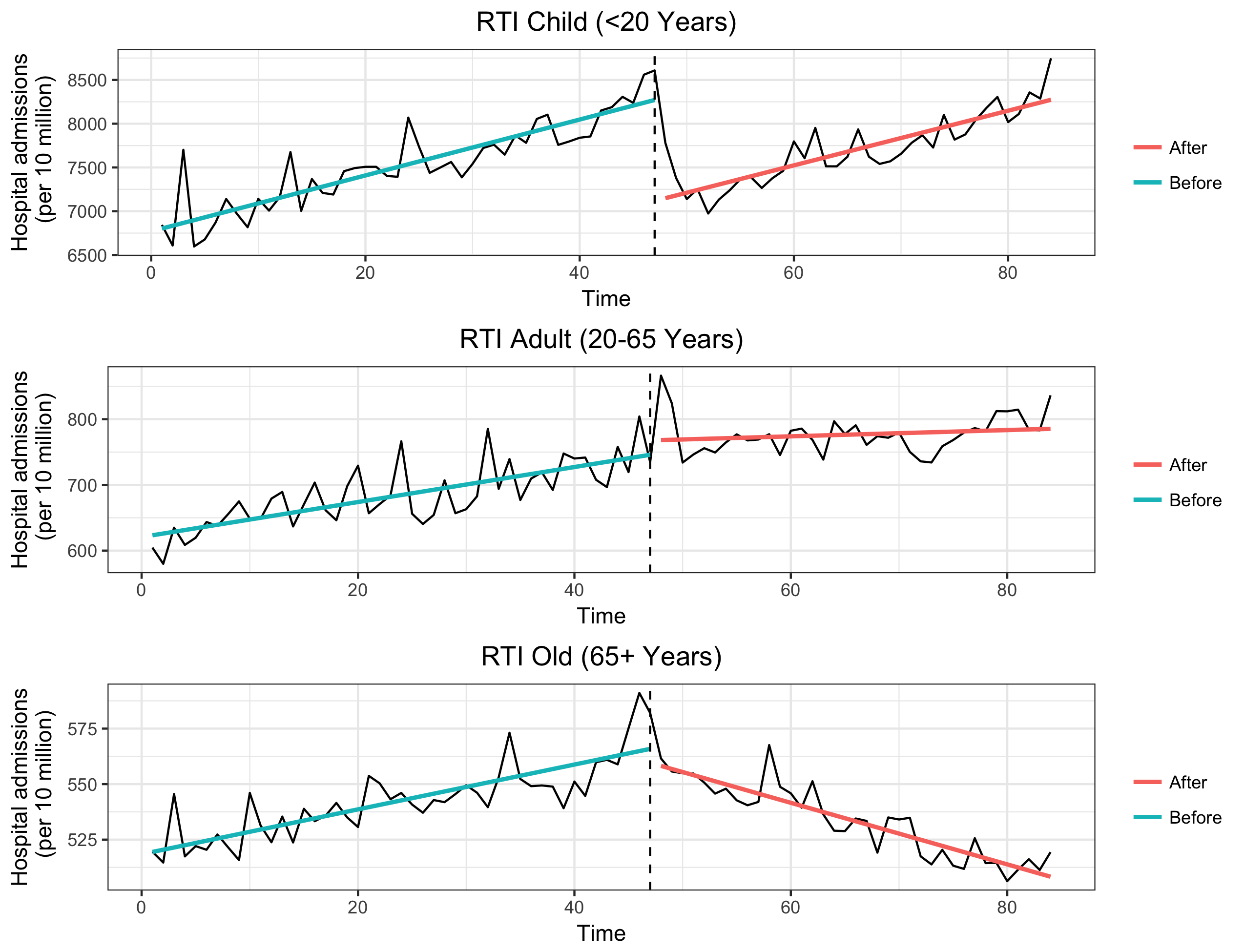

**Fig. 3.** Time series plot of monthly mean age- and sex-standardised emergency hospital admissions: adjusted for; seasonality, air pollutants and time trend, for RTI diseases in Hong Kong, 2010-2016.

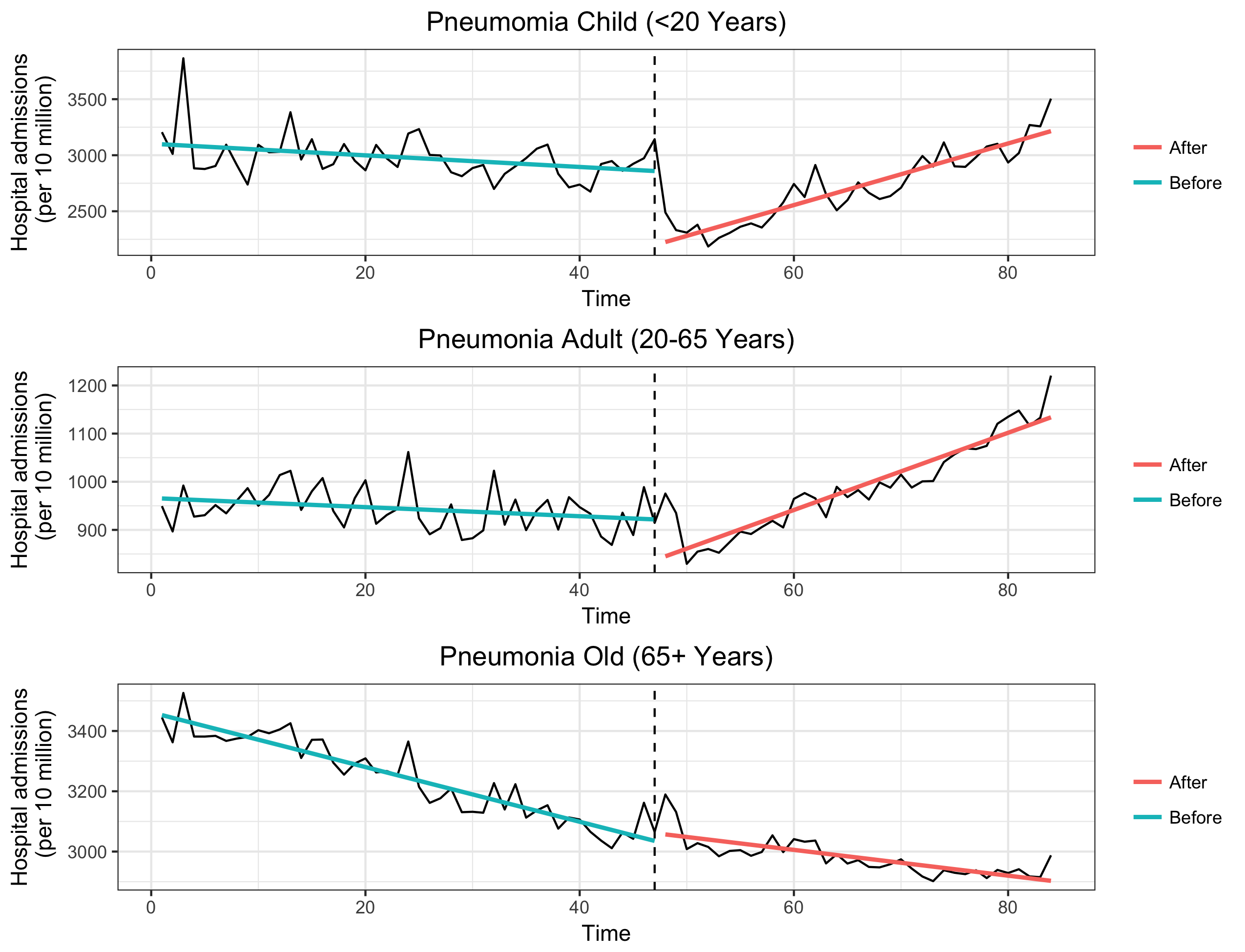

**Fig. 4.** Time series plot of monthly mean age- and sex-standardised emergency hospital admissions, adjusted for; seasonality, air pollutants and time trend, for Pneumonia diseases in Hong Kong, 2010-2016.

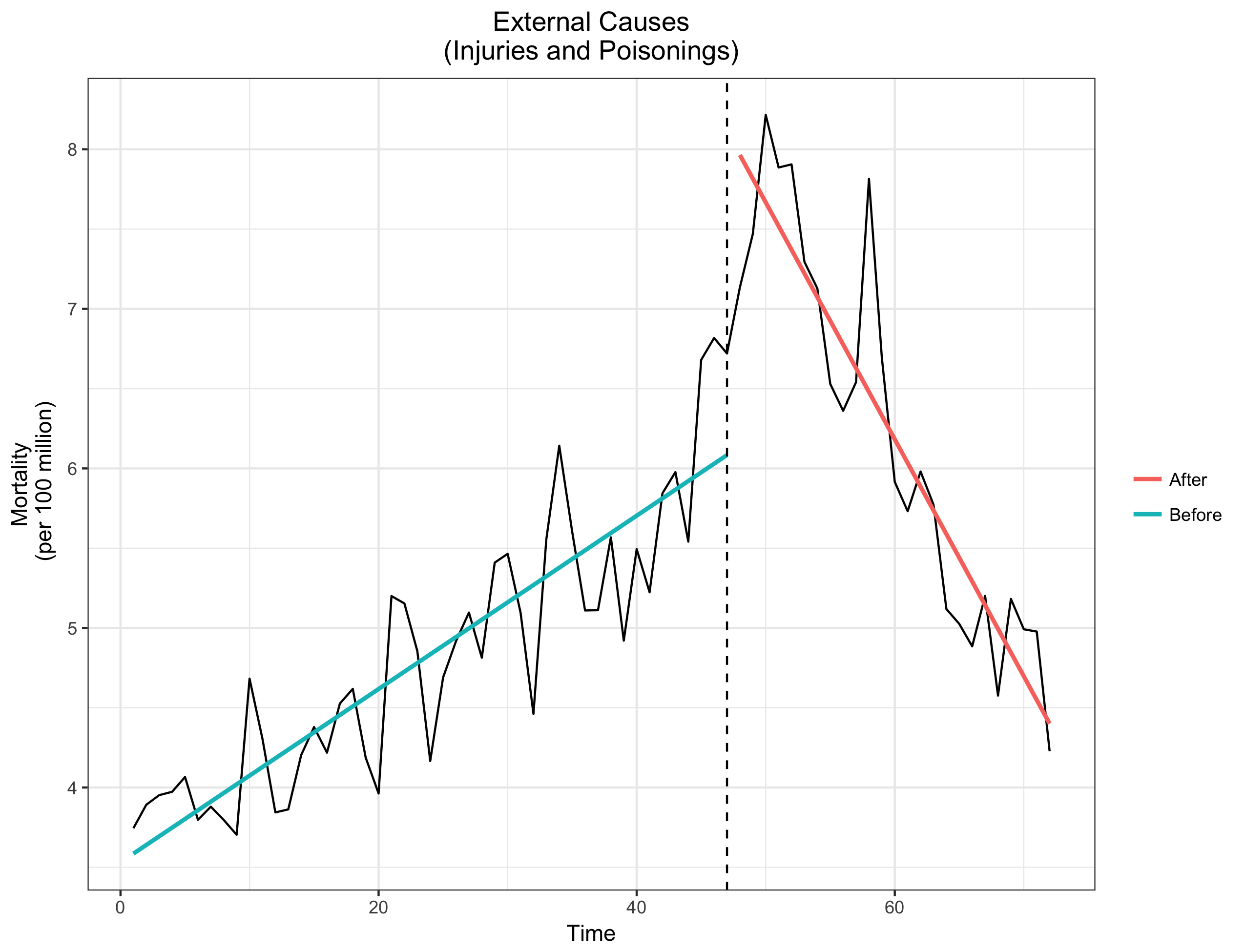

**Fig. 5**. Time series plot of monthly mean, age- and sex-standardised mortality: adjusted for; seasonality, air pollutants and time trend, for injuries, poisonings and certain other consequences of external causes in Hong Kong, 2010-2016.

**Supplementary**

**Table 5** Approximation of immediate and gradual changes in respiratory diseases, at three false policy periods (3 months, 6 months and 12 months) before the original AQHI policy date†.

| Diseases | Immediate Effects- RR (95% CI) | Gradual Effects- RR (95% CI) |
| --- | --- | --- |
| **3 months** |  |  |
| All Respiratory diseases | 0.95 (0.83-1.08) | 1.00 (1.00-1.01) |
| Respiratory tract infection | 0.89 (0.78-1.01) | 1.00 (0.99-1.00) |
| Asthma | 0.98 (0.86-1.12 | 1.00 (0.99-1.01) |
| COPD | 0.99 (0.91-1.07) | 1.00 (1.00-1.01) |
| Pneumonia | 0.92 (0.80-1.07) | 1.00 (1.00-1.01) |
| **6 months** |  |  |
| All Respiratory diseases | 1.00 (0.88-1.14) | 1.00 (1.00 -1.01) |
| Respiratory tract infection | 0.96 (0.88-1.10) | 1.00 (0.99-1.00) |
| Asthma | 0.97 (0.85 -1.10) | 1.00 (0.99-1.01) |
| COPD | 1.00 (0.92-1.08) | 1.00 (1.00-1.01) |
| Pneumonia | 1.01(0.88-1.17) | 1.01 (1.00-1.02) |
| **12 months** |  |  |
| All respiratory diseases | 1.08 (0.94-1.24) | 1.00 (1.00-1.01) |
| Respiratory tract infection | 0.93 (0.82-1.06 | 1.00 (0.99-1.01) |
| Asthma | 0.98 (0.87-1.11) | 1.00 (0.99-1.01) |
| COPD | 0.90 (0.83-0.97) | 1.01 (1.00-1.01) |
| Pneumonia | 0.93(0.82-1.06) | 1.00 (0.99-1.01) |

†Adjusted for seasonality, time trend, SO_2_, NO_2_, PM_10_, O_3._
